# Comparative analysis of macroalgae supplementation on the rumen microbial community: *Asparagopsis taxiformis* inhibits major ruminal methanogenic, fibrolytic, and volatile fatty acid-producing microbes *in vitro*

**DOI:** 10.1101/2022.09.08.507231

**Authors:** E O’Hara, P Moote, S Terry, KA Beauchemin, TA McAllister, DW Abbott, RJ Gruninger

## Abstract

Seaweeds have received a great deal of attention recently for their potential as methane-suppressing feed additives in ruminants. To date, *Asparagopsis taxiformis* has proven a potent enteric methane inhibitor, but it is a priority to identify local seaweed varieties that may hold similar properties. It is essential that any methane inhibitor does not compromise the function of the rumen microbiome. In this study, we conducted an *in vitro* experiment using the RUSITEC system to evaluate the impact of *A. taxiformis, Palmaria mollis*, and *Mazzaella japonica* on rumen prokaryotic communities. 16S rRNA sequencing showed that *A. taxiformis* had a profound effect on the microbiome, particularly on methanogens. Weighted Unifrac distances showed significant separation of *A. taxiformis* samples from the control and other seaweeds (P<0.05). Neither *P. mollis* nor *M. japonica* had a substantial effect on the microbiome (P>0.05). *A. taxiformis* reduced the abundance of all major archaeal species (P<0.05), leading to an almost total disappearance of the methanogens. Prominent fibre-degrading and volatile fatty acid (VFA)-producing bacteria including *Fibrobacter* and *Ruminococcus* were also inhibited by *A. taxiformis* (P<0.05), as were other genera involved in propionate production. However, the abundance of many other major bacteria (e.g. *Prevotella*) was increased by *A. taxiformis* suggesting the rumen microbiome adapted to an initial perturbation. Our study provides baseline knowledge of microbial dynamics in response to seaweed feeding over an extended period and suggests that feeding *A. taxiformis* to cattle to reduce methane may directly or indirectly inhibit important fibre-degrading and VFA-producing bacteria.

## 2. Introduction

To combat the ongoing climate change crises, international legislative agreements have mandated limits on global warming to 1.5°C and 2°C by 2030 and 2050, respectively (Arndt et al., 2022), targets which are unlikely to be met if global food systems continue business as usual practices (Clark et al., 2020). The need for the adoption of environmentally sustainable practices is tempered by the growing nutritional requirements of a worldwide population expected to reach 9.5 bn by 2050. Food production industries are significant sources of anthropogenic carbon, contributing around 30% of total greenhouse gas (GHG) emissions annually, with livestock responsible for ∼30% of this total (Arndt et al., 2022). Numerous GHG including carbon dioxide (CO_2_) and nitrous oxide (N_2_O) are produced throughout the food supply chain, but methane (CH_4_) is the most prominent GHG associated with ruminant production. Methane has a 100-year global warming potential 28-times greater than that of CO_2_ (IPCC, 2013), and enteric methanogenesis contributes 88% of the CH_4_ emissions derived from beef and dairy industries (Caro et al., 2016; FAO GLEAM, 2017).

Microbial fermentation in the rumen facilitates the breakdown of host-indigestible plant biomass, meeting up to 70% of the animal’s energy requirements principally via the production of the volatile fatty acids (VFA) acetate, propionate, and butyrate (Bergman, 1990). However, synthesis of acetate and butyrate also generates significant amounts of hydrogen gas (H_2_) which is converted to CH_4_ by methanogenic archaea and released via eructation (Morgavi et al., 2010). Rumen methanogenesis also represents a loss of up to 12% of the gross energy intake of cattle (Johnson & Johnson, 1995), and thus developing effective ruminal CH_4_ mitigation strategies is attractive from both environmental and economic perspectives. Multiple avenues to reduce enteric methanogenesis in ruminants have been explored, including vaccination, selective breeding, dietary manipulation, improvements in feed efficiency, and supplementation with anti-methanogenic compounds (Gerber et al., 2013). The synthetic inhibitor 3-nitrooxypropanol (3-NOP; Bovaer™, DSM Nutritional Products, Basel, Switzerland) has proven particularly effective, reducing CH_4_ production by up 82%, though results vary across diets and species and reductions of between 20-30% are more typical (Alemu et al., 2021; Hristov et al., 2015; Yu et al., 2021).

There is a long history of seaweed use in human (Peñalver et al., 2020) and livestock (Makkar et al., 2016) diets. Seaweeds contain an abundance of health-promoting secondary metabolites and bioactives (e.g. phlorotannins) due to their antioxidant, antimicrobial, and anti-inflammatory properties (Lomartire et al., 2021). Recently, the addition of seaweeds and their byproducts to ruminant diets has proven highly successful in suppressing enteric methanogenesis, presenting a promising and renewable approach to CH_4_ mitigation (Abbott et al., 2020). Several species of red and brown macroalgae are known to inhibit microbial methanogenesis and have been investigated for their potential as an anti-methanogenic feed additive in ruminants (Machado et al., 2014, 2016b, 2018; Roque et al., 2019). In particular, the red seaweed *Asparagopsis taxiformis* possesses potent anti-methanogenic properties both *in vitro* and *in vivo*, capable of reducing CH_4_ production by over 95% (Roque et al., 2019, 2021; Terry et al., 2022), though potentially at the cost of reduced VFA production (Machado et al., 2016a). The anti-methanogenic property of algae like *A. taxiformis* is mainly due to high bromoform content, a halogen known to inhibit methanogenesis (Machado et al., 2016b). However, there are some concerns that feeding seaweed to ruminants may lead to a spike in milk and/or meat bromoform content with potential implications for consumer health (Muizelaar et al., 2021). The inhibitory effect of *A. taxiformis* on CH_4_ production is also greater than that of concentrated halomethanes, indicating that multiple bioactives present in seaweeds may be working synergistically to suppress methanogenesis (Machado et al., 2018). The bioactives in seaweed may confer other benefits beyond CH_4_ reduction such as reducing fecal pathogen shedding (Zhou et al., 2018), thus non anti-methanogenic seaweeds may hold value as feed additives for ruminants.

*A. taxiformis* is native to warm temperate waters and considered invasive elsewhere, and therefore identification of local varieties of anti-methanogenic seaweeds is of great priority. Recently we examined the effects of two red algae, *Mazzaella japonica* and *Palmaria mollis*, on microbial fermentation and CH_4_ production in a simulated rumen environment but found no tangible impact on methanogenesis (Terry et al., 2022). However, it is unknown if these seaweeds might influence the rumen microbiome in other ways and understanding of the microbial mechanisms underpinning ruminal adaptation to seaweed supplementation is generally lacking. To address this, we employed 16S rRNA amplicon sequencing to assess temporal dynamics of the rumen microbiome in response to supplementation with *A. taxiformis, M. japonica*, and *P. mollis* over a 13-day period using a RUSITEC apparatus. These data provide insight into rumen microbial adaptation to seaweed feeding, but also highlight potentially negative impacts on key rumen functions concurrent to CH_4_ inhibition.

## 3. Materials and Methods

### 3.1. RUSITEC System

Details of seaweed procurement, RUSITEC design, and sampling protocols have been published elsewhere (Terry et al., 2022). Briefly, this experiment was performed in duplicate using two identical RUSITEC apparatus equipped with eight fermentation vessels each. The experiment was divided into four phases: baseline (days 0-7), adaptation (days 8-11), intermediate (days 12-16) and stable (days 17-21). Each fermenter was inoculated with rumen fluid collected from 3 ruminally canulated beef heifers. The basal substrate (10 g dry matter (DM) 50:50 barley silage:barley straw) was added daily to the fermenters. The three seaweeds, *Asparagopsis taxiformis, Mazzaella japonica* and *Palmaria mollis* were added to the basal substrate on a 2% DM basis from day 8 onward. The control vessels were treated in an identical manner with no seaweed added. A 1.5 ml sample of digester fluid was collected daily from each vessel for microbial community analysis. The sample was centrifuged at 21,000 RCF for 5 min to pellet the microbial cells. The supernatant was discarded and the cell pellet was snap frozen in liquid nitrogen and stored at -80°C, pending molecular analysis.

### 3.2. Bromoform content analysis

The concentration of bromoform in each of the three seaweeds was determined using a Shimadzu QP2010 Ultra GC/MS system at a commercial laboratory (Bigelow Analytical Services, USA). Bromoform concentration was recorded as mg/g dry weight.

### 3.3. DNA isolation

Microbial DNA was isolated from all samples using the ZymoBIOMICS DNA Miniprep Kit, following the manufacturer’s instructions with some modifications; the cell pellet was suspended in 1000 μl of DNA/RNA Shield (Zymo Research) and 250 μl of this cell suspension was added to 250 μl of lysis buffer in ZR BashingBead™ lysis tubes (Zymo Research) containing 0.1 mm and 0.5 mm beads. All samples underwent 5 rounds of bead beating on a MP FastPrep 24 (MP Bio, USA) at 6.0 m/s with 20 s breaks between rounds. Following bead beating, samples were incubated at 70°C for 10 min to increase the effectiveness of DNA isolation. The remaining purification steps followed the manufacturers protocol, and all samples were eluted in 100 μl of DNA/RNA Shield. DNA quantity and quality was assessed by 2 consecutive readings on a Nanodrop2100 spectrophotometer.

### 3.4. PCR amplification, library construction, and sequencing

The V4 hypervariable region of the 16S rRNA gene was amplified with the primer pair 515F (5′-GTGCCAGCMGCCGCGGTAA-3′) and 806R (5′-GGACTACHVGGGTWTCTAAT-3) on a Fast Start High Fidelity PCR System (Roche, PQ) using the following conditions: 94°C for 2 min, followed by 33 cycles of 94°C for 30 s, 58°C for 30 s and 72°C for 30 s, with a final elongation at 72°C for 2 min. Libraries were prepared using the Illumina MiSeq Reagent Kit V2 and sequenced on an Illumina MiSeq using 2×250 bp chemistry at a commercial sequencing laboratory (Genome Quebec, QC, Canada).

### 3.5. Sequence data analysis

Raw reads were imported in .fastq format to a local server for analysis. Read quality was evaluated using FASTQC (Andrews, 2010) and MultiQC (Ewels et al., 2016). Data were processed using the QIIME2 software package (Bolyen et al., 2019). FIGARO (Weinstein et al., 2019) was used to identify optimum truncation positions for read merging. Reads were denoised into amplicon sequencing variants (ASVs) using the DADA2 (Callahan et al., 2016) plugin for QIIME2. A phylogenetic tree was generated using MAFFT (Katoh & Standley, 2013). ASVs were taxonomically classified using a Native Bayes classifier trained on the V4 region of the 16S rRNA gene using the Sci-Kit plugin in QIIME2. The SILVA SSU (v.138.1) (Quast et al., 2013) database was used to classify bacterial sequences while the Rumen and Intestinal Methanogens (RIM) database (v.14.7) (Seedorf et al., 2014) was used for archaeal reads. Initial analysis (described in Supplementary Data) indicated that increasing the confidence threshold to 0.85 (from a default of 0.7) prevented spurious ASV classification using the RIM database.

### 3.6. Diversity and Correlation analysis

QIIME2 objects (frequency table, taxonomy table, phylogenetic tree) were imported into R as a Phyloseq (McMurdie & Holmes, 2013) object using the qiime2R package (Bisanz, 2018). Analysis was performed separately for bacterial and archaeal datasets. In-house R scripts were used to calculate summary statistics of read counts and distributions. ASVs that were not assigned to at least a microbial phylum were discarded. Rarefaction curves were generated to identify suitable subsampling thresholds for diversity analyses. Rarified data was used for ɑ-(Chao1 and Shannon) and β-diversity (Weighted Unifrac) calculations. ɑ-Diversity data was summarized according to phase for comparison using a two-way analysis of variance (ANOVA) with Tukeys post-hoc test. Statistically significant differences were declared at Bonferroni-adjusted P<0.05. Permutational Analysis of Variance (PERMANOVA) tests were performed using the Vegan (Oksanen et al., 2020) R package, with pairwise tests performed using the RVaideMomoire R package (Herve, 2021).

Homogeneity of Dispersion (β-dispersion) tests were performed using Vegan. Canonical Correspondence Analysis (CCA) between rarified ASV count data and environmental data collected from the RUSITEC system was also performed using Vegan. Statistically significant differences were declared at a threshold of P<0.05, and all figures were generated using ggplot2 in R (Wickham, 2009). Core rumen taxa were declared as those present in more than 50% of the samples and represented by at least 100 sequencing reads. Correlation analysis was performed between selected fermentation metrics and (i) the core microbiome and (ii) differentially abundant taxa using log10 transformed count data and Spearman correlations in R.

### 3.7. Differential abundance testing

Differentially abundant (DA) taxa between seaweed and control samples were identified using ANCOM-BC and ALDEx2. Data were arranged according to phase (adaptation: days 8-11; intermediate: days 12-15; stable: days 16-19). Prior to DA testing, low-prevalence features - those present in <10% of the samples - were discarded as recommended for microbiome data (Nearing et al., 2021). Non-rarefied data were used as input for all DA tests. For ANCOM-BC, the phyloseq object was passed to the ‘ancombc’ function and run at a maximum of 1000 iterations. Structural zeros - taxa present in one group but absent in another - were declared as DA where present. A pseudocount of 1 was added to all observations to facilitate log transformation. Significant features were those with a Benjamini-Hochberg-adjusted P< 0.05. For ALDEx2 count data and metadata were passed to the ‘aldex’ command which employs a Dirichlet-multinomial model to infer abundance from counts. P-values generated from a Wilcoxon Rank Sum test were FDR-corrected using the Benjamini-Hochberg procedure, and significant differences according to treatment group were declared at corrected P<0.05. The lists of significantly different taxa identified by both tools were compared, with the consensus taxa (i.e., those identified using both methods) declared as DA for each respective comparison.

## 4. Results

### 4.1. Bromoform analysis

Only *A. taxiformis* had detectable amounts of bromoform, at 0.517 mg/g^-1^ DM. Any bromoform present in *P. mollis* and *M. japonica* was below the limit of detection.

### 4.2. Sequencing data characteristics

Sequencing of 16S rRNA V4 amplicons from 272 digesta samples generated a total of 12,456,461 reads (range: 2,663 – 78,773) with an average of 45,795 ± 10,306 (mean ± SD) per sample. Denoising with DADA2 retained 72.58% of the reads and identified a total of 18,925 ASVs. Rarefaction curves for both bacterial and archaeal annotation data reached a plateau (Supplementary Data) indicating that our sampling depth was sufficient.

### 4.3. Effect of seaweed supplementation on microbial communities in the RUSITEC system

#### 4.3.1. Alpha and Beta diversity

Principal coordinate (Figure 1) and PERMANOVA (Table 1) analyses based on weighted Unifrac distances showed separation of both bacterial and archaeal microbial profiles according to treatment and phase (P<0.05), with the *A. taxiformis* samples clearly separating from the other groups. The separation increased throughout the experiment and was greatest in the stable phase (Figure 1a). The R^2^ value was greater for Treatment than Phase (0.30 vs. 0.04) in the bacterial data, indicating this was the largest factor contributing to compositional differences, but Phase and Treatment made similar (0.17 vs. 0.18) contributions to dissimilarity of the archaeal data. Canonical correspondence analysis (Figure 1c) using the fermentation data reported separately by Terry et al (2022) showed that the separation of the *A. taxiformis* samples from the other groups could be explained by several environmental measurements, including the decline in CH_4_, and concordant increase in H_2_ production, as well as the lower molar proportions of propionate and acetate (Terry et al 2022). The environmental parameters that drove the separation of samples was similar for both the bacterial and archaeal datasets (Figure 1d).

**Table 1.** Results of PERMANOVA analysis on bacterial and archaeal community dissimilarities. Tests conducted using a Weighted Unifrac dissimilarity matrix.

| Variable | R<sup>2</sup> | F-value | P-value | R<sup>2</sup> | F-value | P-value |
| --- | --- | --- | --- | --- | --- | --- |
|  | Bacteria |  |  | Archaea |  |  |
| Treatment | 0.30 | 35.63 | 0.001 | 0.18 | 20.02 | 0.001 |
| Phase | 0.05 | 8.41 | 0.001 | 0.17 | 28.40 | 0.001 |
| Day | 0.01 | 2.18 | 0.065 | 0.01 | 2.60 | 0.064 |
| Treatment:Day | 0.07 | 7.75 | 0.001 | 0.02 | 2.25 | 0.025 |
| Treatment:Phase | 0.02 | 1.03 | 0.416 | 0.03 | 1.44 | 0.124 |
| Residual | 0.56 |  |  | 0.60 |  |  |
| Total | 1.00 |  |  | 1.00 |  |  |

\*R<sup>2</sup>: Proportion of variance explained by each variable; F-value: estimate of effect size.

**Figure 1:**
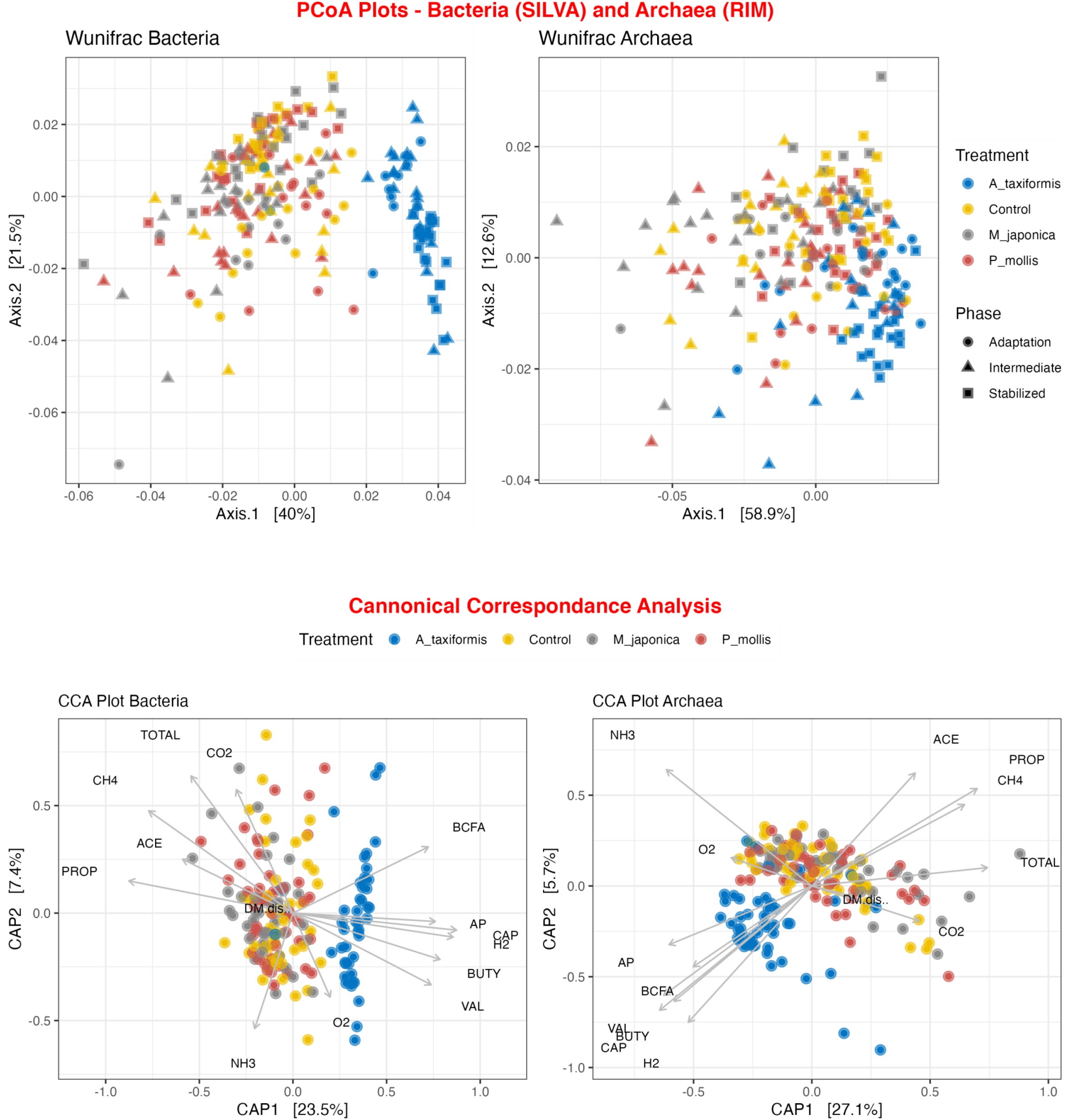
Principal Coordinate Analysis (PCoA) plots of (a) bacterial and (b) archaeal communities. Canonical correspondence (CCA) plots of (c) bacterial and (d) archaeal communities and fermentation variables. All plots were generated from a weighted Unifrac distance matrix.

ANOVA analysis indicated that Chao1 and Shannon indices were influenced by treatment (P<0.05), phase (P<0.05), and the interaction of the two (P<0.05). Only *A. taxiformis* exhibited major differences in alpha-diversity when compared to the controls (Figure 2), while differences between the controls and the other 2 seaweeds were minor. All comparisons are listed in supplementary information. None of the seaweed treatments caused temporal changes in diversity or evenness between phases (P>0.05). *A. taxiformis* samples had lower Chao1 and Shannon index values in the intermediate phase compared to the controls (P<0.05), while the *P. mollis, M. japonica* and Control samples had similar values. Treatment-wise differences declined for all treatments by the Stable phase with few statistically significant differences evident.

**Figure 2:**
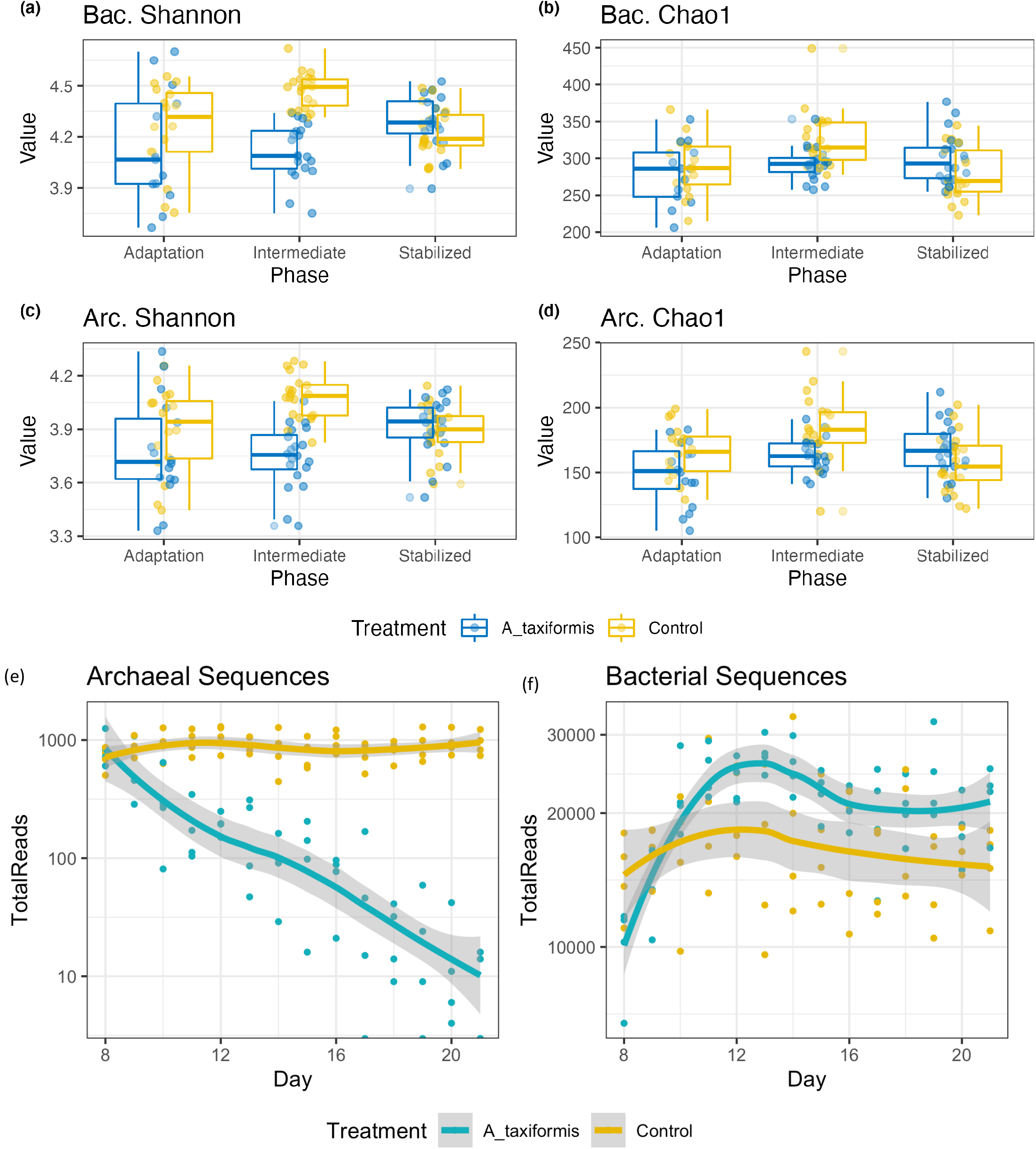
Alpha diversity boxplots of bacterial and archaeal sequencing data. Samples are grouped according to phase. All data was rarefied to an even depth prior to metric calculation. Panels (a-d) depict diversity metrics, panels (e) and (f) show total archaeal sequences respectively.

#### 4.3.2. Microbial community composition and response to seaweed addition

Following the removal of sparse ASVs, 1 archaeal and 16 bacterial phyla were identified across all samples. Irrespective of treatment or phase, the microbiomes were dominated by Bacteroidota and Firmicutes throughout, with Proteobacteria and Actinobacteria also present at high proportions. The most prominent rumen bacteria included *Prevotella, Streptococcus, Megaspheara* and *Lactobacillus* (Figure 3). The “core” bacteriome was calculated across Control and *A. taxiformis* samples and consisted of 32 genera drawn from 8 phyla. The divergence in the core microbial groups between Control and *A. taxiformis* samples increased across phases (Supplementary Data) indicating that at least some of these taxa were influenced by *A. taxiformis* supplementation in the RUSITEC. As expected, major ruminal bacteria including *Fibrobacter, Prevotella, Succinivibrio, Megasphaera*, and *Butyrivibrio* were part of the core microbiome. The methanogen community was dominated by *Methanobrevibacter ruminantium, Methanobrevibacter gottschalkii*, and several poorly characterized species belonging to *Methanomassilicoccaceae* (Figure 4). *Methanomicrobium mobile, Methanimicrococcus blatticola*, and *Methanosphera* sp. were among the less abundant methanogens in the RUSITEC. When raw abundances were visualized, there was a clear decline in the number of sequences attributed to methanogenic species over time in the *A. taxiformis* samples (Figure 4).

**Figure 3:**
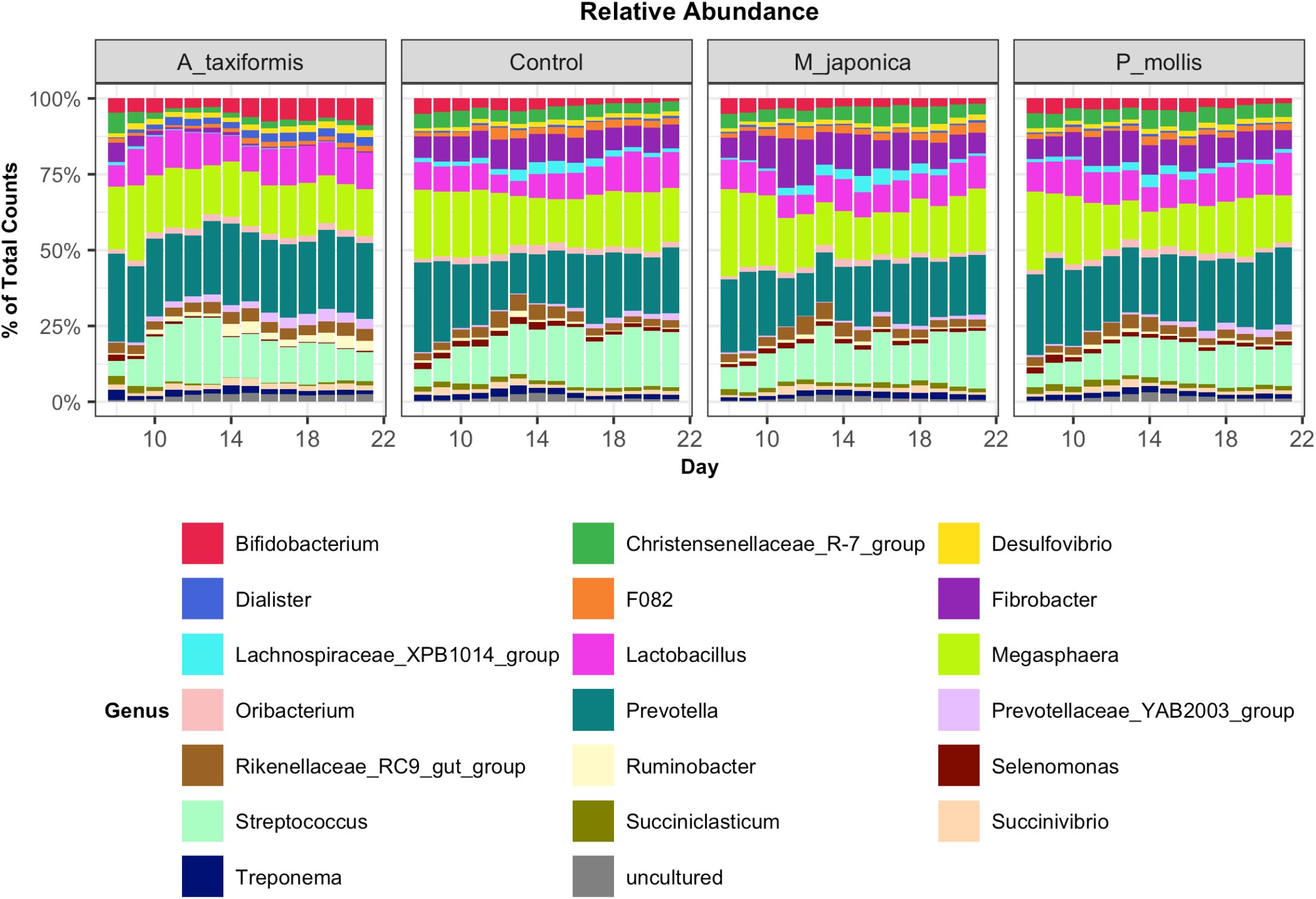
Relative abundances of the 20 most abundant genus-level features across all treatment groups. Abundances were scaled to 1 for ease of presentation.

**Figure 4:**
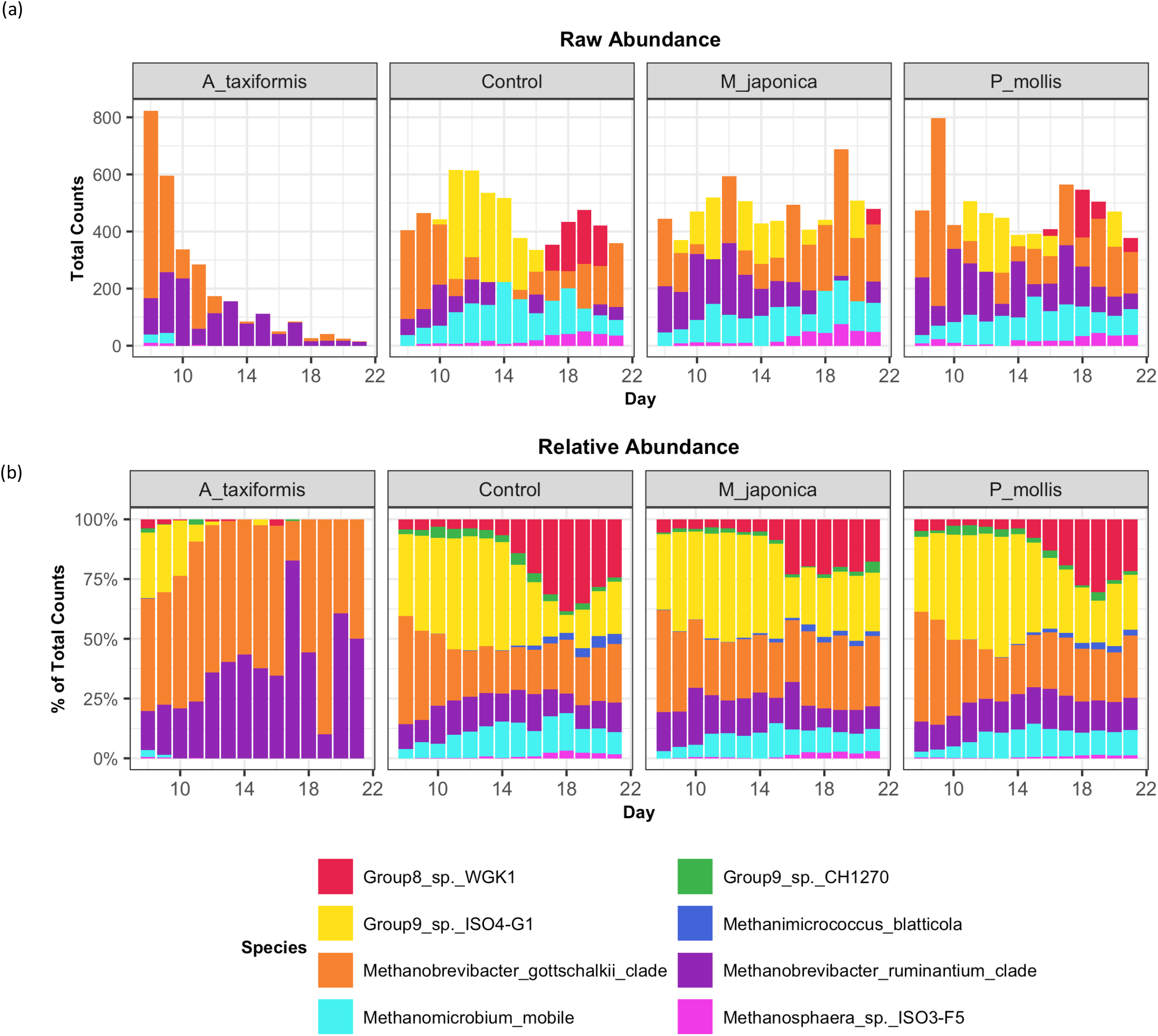
(a) Raw and (b) Relative abundances of methanogenic species across all treatment groups. Values are the daily median across all samples.

The microbial response underpinning the dramatic reduction in CH_4_ by *A. taxiformis* reported by Terry et al. (2022) was investigated in more detail via correlation analysis and differential abundance testing. To examine if the bioactives present in non-methanogenic seaweeds had an impact on the microbiome, testing was also conducted between the control and *M. japonica* samples. Differentially abundant (DA) features were those identified using both tools (see methods). *M. japonica* did not have a major effect on the bacterial community, with just 3 genera responding over the course of the experiment; during the adaptation phase, a genus-level feature from the family *Selenomonadaceae* was more abundant in the seaweed samples than in the controls (P<0.05). There was no impact of *M. japonica* during the intermediate phase, while during the adaption phase, two DA genera were identified: *Oribacterium* and *Pirellulaceae P1088_a5_gut_group* (P<0.05).

The bacterial community exhibited a progressive response to *A. taxiformis*, with 10, 76, and 92 DA bacterial genera identified during the adaptation, intermediate, and stable phases, respectively (P<0.05). ANCOM-BC identified more DA genera throughout the experiment than Aldex2 (Figure 5a). Nine genus-level features were affected by *A. taxiformis* during all 3 phases. Among them, the proportions of *Fibrobacter, Schwartzia*, and *Papillibacter* were reduced by *A. taxiformis* throughout the experiment compared to the controls (P<0.05), while the abundance of *Sutterella* was the only genus that was consistently higher for *A. taxiformis* compared to the controls (P<0.05). There was a large degree of overlap between the DA taxa in the intermediate and stable phases, with 54 showing the same response to seaweed supplementation in both phases (Figure 5b & 5c). Many major rumen bacteria and members of the core microbiome were influenced by *A. taxiformis* in the latter phases of the experiment, including increased abundance of *Prevotella, Dialister, Succinivibrio* and *Ruminobacter* (P<0.05), while *Clostridium sensu stricto 1, Roseburia, and Ruminoclostridium* were among the genera more abundant in the control samples (P<0.05). Moreover, many less noted members of the assemblage responded to *A. taxiformis* in the intermediate and stable phases, including lowered abundances of *Endomicrobium, Denitrobacterium, and Angelakisella* (P<0.05). There were also shifts in the abundances of many unclassified and/or poorly annotated genera belonging to *Prevotellaceae* and *Lachnospiraceae* (P<0.05), as well as other families.

**Figure 5:**
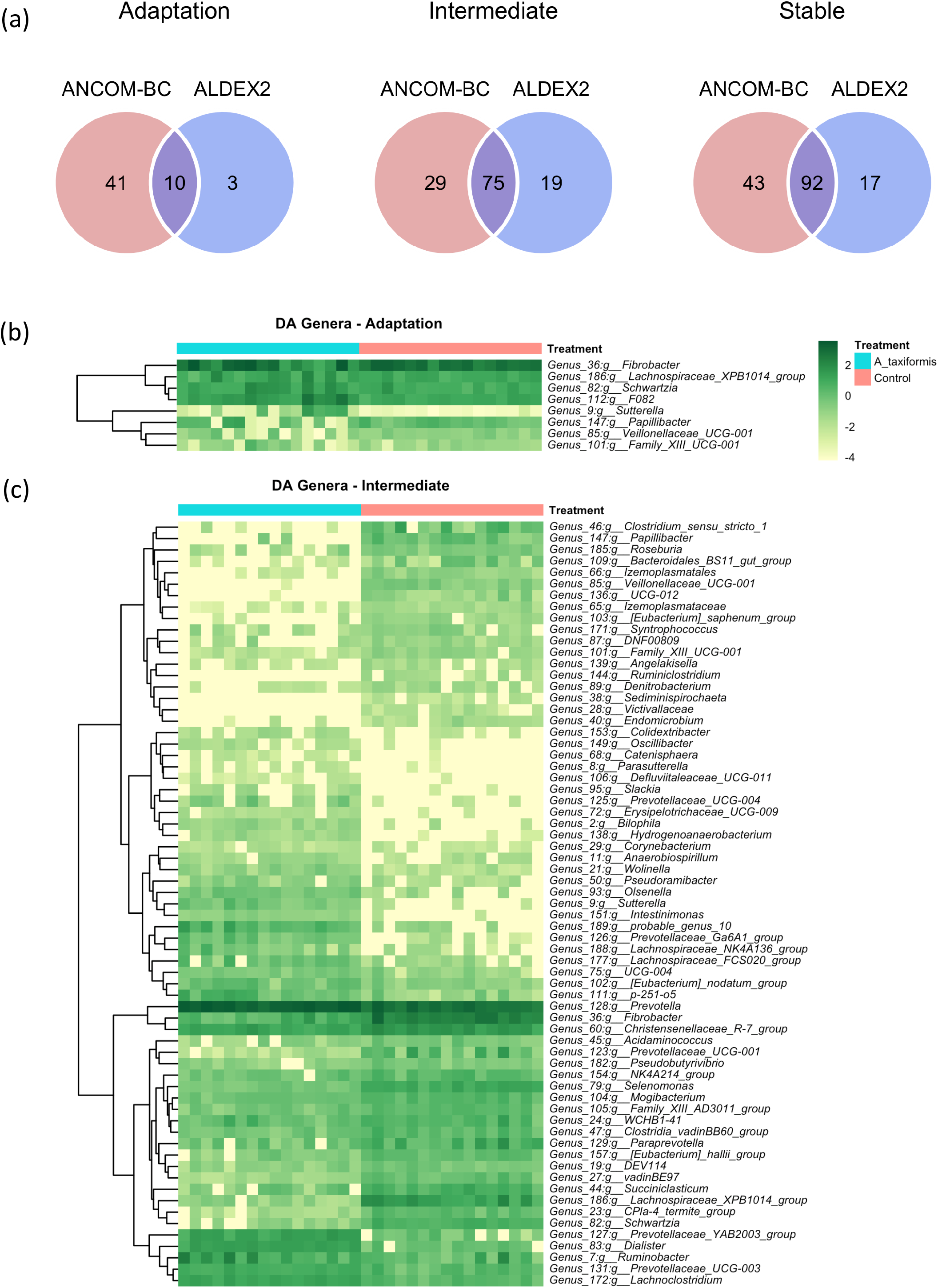
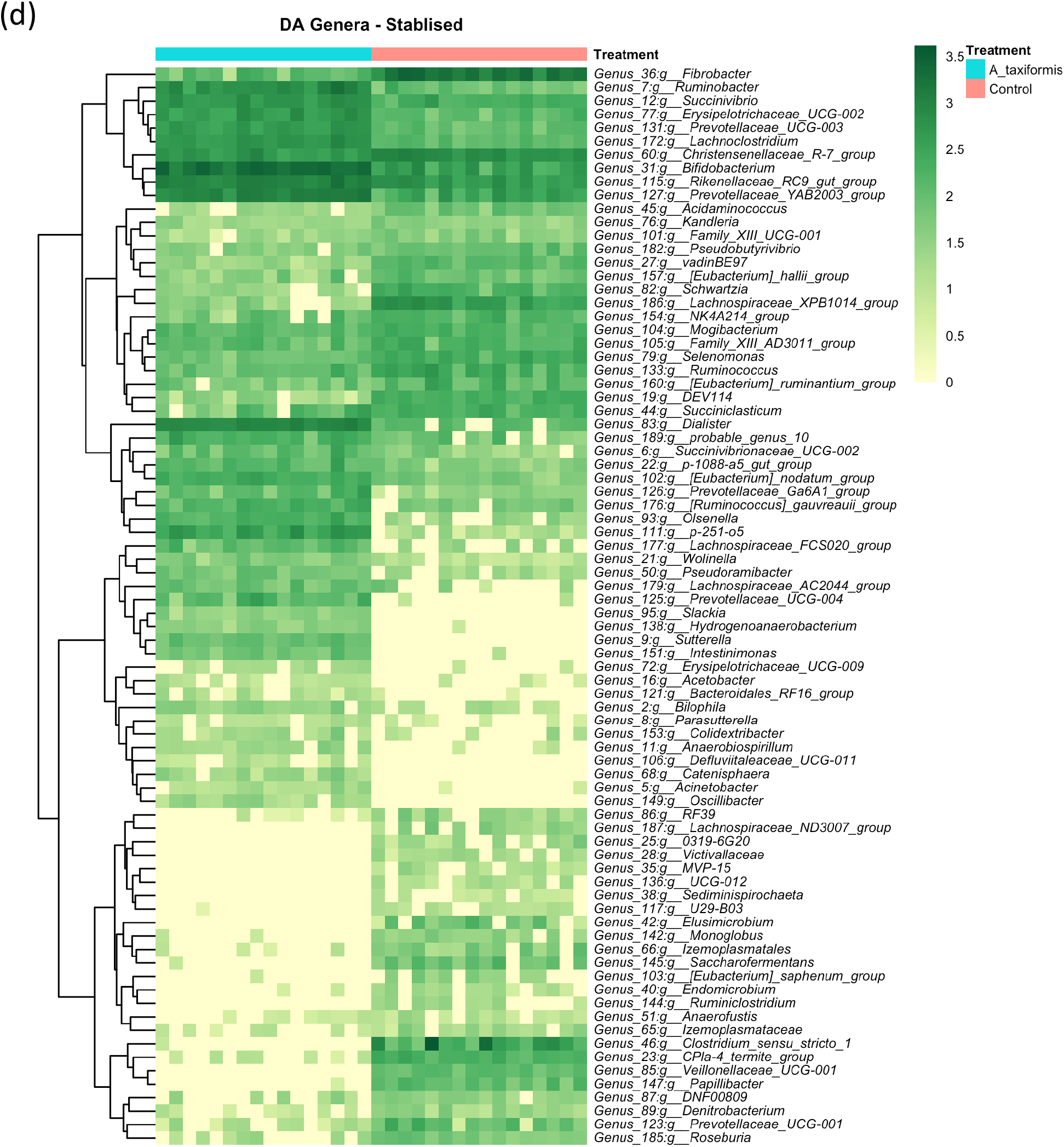
Differential abundance (DA) analysis of bacterial genera. (a) Venn diagrams showing the overlap between ANCOM-BC and Aldex2 results. Heatmaps of DA genera for the (a) adaptation, (b) intermediate and (c) stabilized phase. Raw count data was log transformed for plotting. Genera denoted as uncultured are exclude

When low-prevalence features were removed, there were 8 archaeal species-level taxa left in our dataset. No archaeal species exhibited a statistically significant response to *M. japonica* supplementation throughout the experiment (P>0.05; Supplementary information). The methanogen community exhibited a dramatic response to *A. taxiformis*; 3,396 reads were assigned to archaeal ASVs on D8 of the filtered dataset, declining to just 30 reads by day 21. In contrast, the number of archaeal sequences recovered from the control samples remained relatively stable throughout (Figure 2e). Abundances of *Methanobrevibacter gottschalkii* and *Methanobrevibacter ruminantium* were lower with *A. taxiformis* compared to the controls (P<0.05) in both the adaptation and intermediate phases (Figure 6). *Methanomicrococcus blatticola* was also less abundant with *A. taxiformis* during the intermediate phase. Only *Methanomassillicoccaceae Group 8 sp. WGK1* was differentially abundant (P < 0.05) between the groups during the stable phase. We interpreted the results of differential abundance testing between *A. taxiformis* and control samples with caution due to the very small number of reads recovered from the *A. taxiformis* samples collected during the latter stages of the experiment.

**Figure 6:**
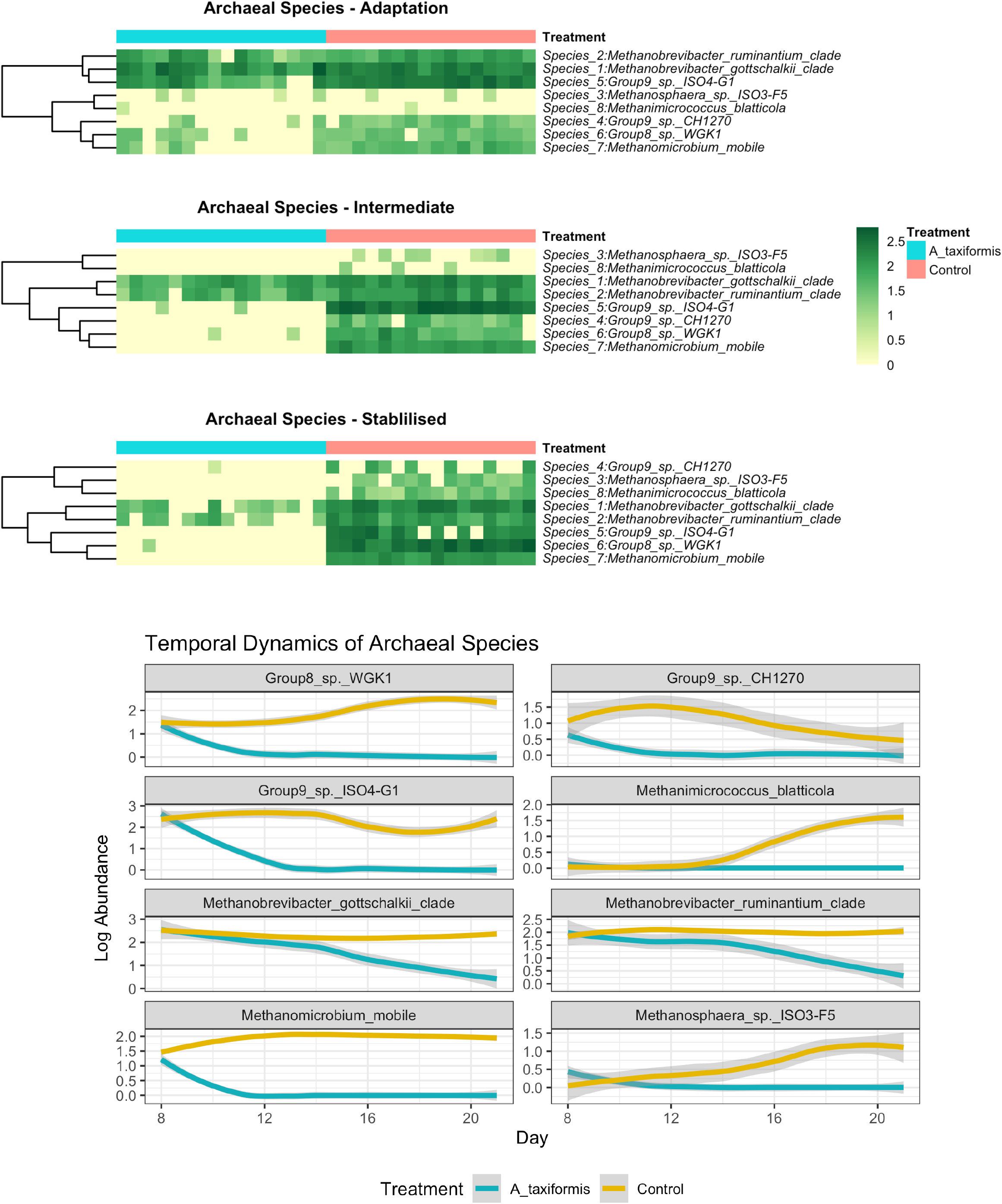
(a) Heatmaps depicting archaeal community composition in each phase and (b) line plots showing temporal species dynamics throughout the experiment. Raw abundances were log-transformed prior to plotting.

#### 4.3.3. Correlation between microbial communities and fermentation variables

Spearman correlation coefficients were used to determine the relationships between fermentation variables reported by Terry and colleagues (2022) and microbiome features. A relationship was considered strong at an absolute R value > 0.5 and an adjusted P-value < 0.05. The core microbes exhibited stronger relationships with fermentation variables in the *A. taxiformis* samples than in the controls. CH_4_ concentration was strongly associated with 13 of the 32 core genera in the *A. taxiformis* samples; *Fibrobacter, Selenomonas, Schwartzia, NK4A214* (*Ruminococcaceae*), and *Lachnospiraceae XPB1014* were all positively correlated with CH_4_ concentration (P<0.05; Figure 7), while *Dsulfovibrio, Ruminobacter, Fibrobacter, Erysipelotrichaceae* UCG-002 and *Dialister* were among those that exhibited negative relationships with CH_4_ (P<0.5). Conversely, only 2 genera correlated with CH_4_ in the control samples, with *Prevotellaceae YAB2003* exhibiting a negative relationship and *Paraprevotella* a positive one (P<0.05). Strong relationships between core taxon abundances and the molar proportions of individual VFA were also more evident in *A. taxiformis* than controls; *Schwartzia* and *Lachnospiraceae XPB1014* group were positively correlated with acetate and propionate (P<0.05), while *Erysipelotrichaceae UCG-002* was negatively correlated with both acids (P<0.05). Proportions of propionate were also negatively correlated with the abundances of *Lactobacillus, Prevotellaceae YAB3003 group*, and *F082* (P<0.05), while *Dialister* and *Prevotellaceae UCG-003* had a negative relationship with acetate proportion (P<0.05). Butyrate was positively associated with *Streptococcus, Megasphaera* and *Oribacterium* in the *A. taxiformis* samples (P<0.05). Total VFA concentration was also negatively correlated with *Desulfovibrio* and *Lactobacillus* in the *A. taxiformis* samples (P<0.05). The proportion of propionate was positively correlated with *Lachnospiraceae XPB1014* in the control samples (P<0.05). Total VFA was positively correlated with *WCHB1-41* and *Paraprevotella* in the controls (P<0.05), while *Erysipelotrichaceae UCG-002* and *Prevotellaceae YAB2003* group exhibited the opposite relationship. Correlation coefficients and P-values for the core genera are provided in the supplementary data and presented graphically in Figure 7.

**Figure 7:**
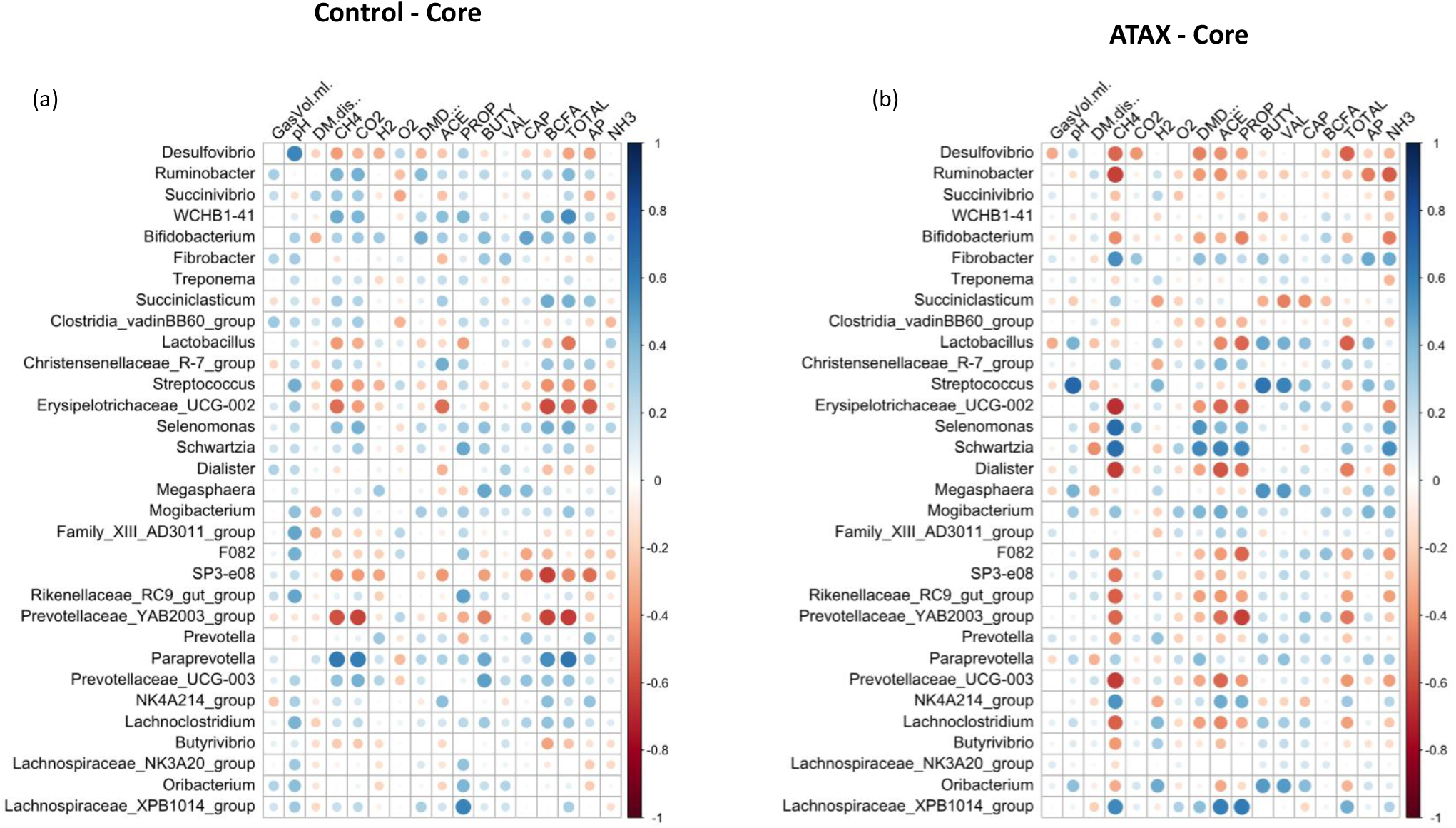
Spearman correlation analysis plots between the core bacteriome and fermentation variables. Raw bacterial abundances were log-transformed prior to correlation. Only data from control and *A. taxiformis* reactors was analyzed.. The size of the circle corresponds to the absolute correlation value. DM.diss = dry matter disappearance; DMD = dry matter digestability; ACE = acetate; PROP = propionate;BUTY = butyrate; VAL = valerate; CAP = caproate; BCFA = branched chain fatty acids; TOTAL = total VFA; A:P = acetate: propionate ratio; NH3 = ammonia

Strong relationships were evident between the DA genera (*A. taxiformis* vs. control) and fermentation variables throughout and largely reflected the DA results. In the adaptation phase, CH_4_ and propionate were positively correlated with multiple DA genera including *Fibrobacter, Schwartzia, Veillonellaceae* UCG-001 and *Lachnospiraceae XPB1014 group* (P<0.05), while *Sutterella* had inverse relationships with both (P<0.05). Conversely, caproate was negatively correlated with 5 of the DA genera, and positively correlated only with *Sutterella* (P<0.05) (Figure 8a). Figure 8b and 8c show that in the latter phases of the experiment, almost all the differentially abundant taxa were significantly correlated with CH_4_ concentration and the molar proportions of VFA (P<0.05). Total VFA, H_2_ concentration, and the acetate:propionate ratio also exhibited strong relationships with DA taxa in the intermediate and adaptation phases (P<0.05).

**Figure 8:**
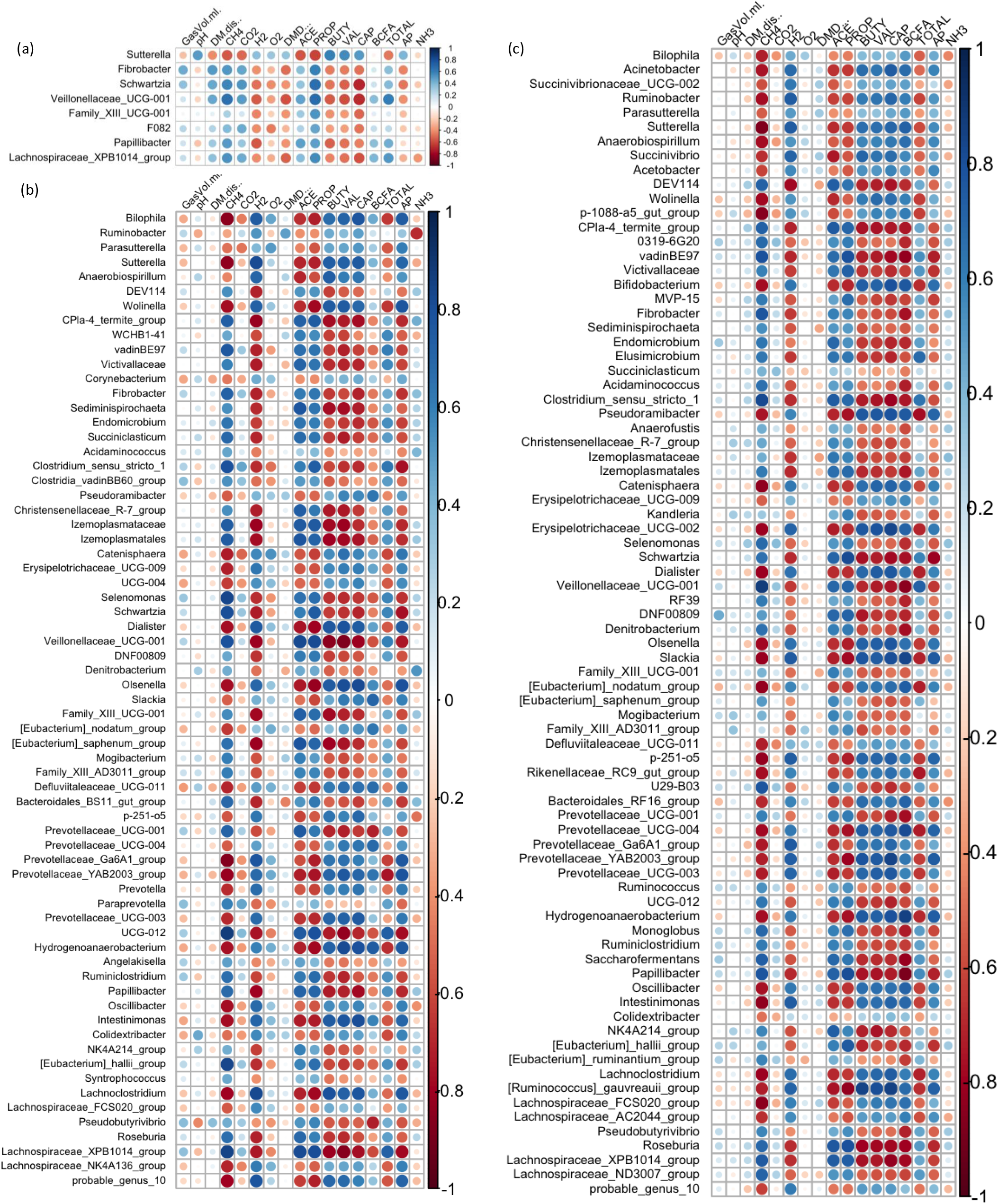
Spearman correlation analysis plots between the differentially abundant genera and fermentation variables during the (a) adaptation, (b) intermediate, and (c) stabilized phases. Raw bacterial abundances were log-transformed prior to correlation. The size of the circle corresponds to the absolute correlation value.DM.diss = dry matter disappearance; DMD = dry matter digestability; ACE = acetate; PROP = propionate;BUTY = butyrate; VAL = valerate; CAP = caproate; BCFA = branched chain fatty acids; TOTAL = total VFA; A:P = acetate: propionate ratio; NH3 = ammonia

## 5. Discussion

The effectiveness of the red seaweed *Asparagopsis taxiformis* in suppressing enteric methanogenesis in ruminants has been demonstrated both *in vitro* and *in vivo* (Kinley et al., 2016; Machado et al., 2016a; Roque et al., 2019, 2021). Our recent study examining the effects of *A. taxiformis, Mazzaella japonica*, and *Palmaria mollis* on *in vitro* rumen fermentation and gas production (Terry et al., 2022) confirmed previous observations of the potency of *A. taxiformis* in reducing methanogenesis, with CH_4_ concentrations declining by 95.1% in the stable phase. However, this mitigation effect was accompanied by reductions in fibre degradation and VFA production, with potential negative implications for animal performance. There was no measurable impact of either *M. japonica* or *P. mollis* on methanogenesis or microbial fermentation. There is limited data concerning the microbial response to seaweed supplementation in ruminants, and this study presented an opportunity to examine the rumen microbial response to seaweed supplementation over an extended period and assess if the microbiomes show any evidence of adaptation to seaweeds which might lessen their anti-methanogenic effects over time as has been previously documented (Knight et al., 2011).

Among the three seaweeds studied, only *A. taxiformis* had a measurable effect on the rumen microbes, reflecting the fermentation data from this experiment that was published in our companion study (Terry et al., 2022). Our analysis of the archaeal reads provided some surprising results, with a high proportion of the reads mapping to *Methanocaldococcus villosus* (described in the supplementary material), a marine archaeon native to sub-marine hydrothermal systems (Bellack et al., 2011) and not known to have any role in ruminal fermentation. Further investigation revealed low classification confidence values (<0.75) of the *M. villosus* ASVs and increasing the confidence threshold from the QIIME2 default of 0.7 to 0.85 eliminated this spurious identification. As a result, we recommend that an ASV assignment confidence threshold of at least 0.85 be employed when utilizing the RIM database for methanogen classification.

The near-total collapse of the methanogen community in the *A. taxiformis* samples was striking, and cannot be attributed to temporal shifts in community composition commonly associated with RUSITEC fermenter apparatus (Mateos et al., 2017) or the decline in protozoa associated methanogens (Roque et al., 2019) as the archaeome of the control samples remained relatively stable throughout. Many sparsely detected archaeal ASVs were present in the dataset, and when these were filtered out for differential abundance testing as recommended (Nearing et al., 2022) the stark differences in archaeal composition following *A. taxiformis* addition became more evident. There was no suggestion of niche transition among the methanogen species following *A. taxiformis* addition, with the abundances of all major archaea declining throughout the experiment, and CCA analysis was suggestive of a strong relationship between the CH_4_ and H_2_ concentrations reported previously (Terry et al., 2022) and methanogen community dynamics. The extent of the decline in methanogen abundance observed here has not, to our knowledge, been previously documented. A recent *in vitro* study showed that while *A. taxiformis* decreased the abundance of methanogenic groups over a 96 h period, the reduction was not as dramatic as we observed (Roque et al., 2019).

Interestingly, in our study *Methanomassilicoccaceae* species declined in abundance almost immediately (within 24 h) after *A. taxiformis* addition, while it took several days for the *Methanobrevibacter* species to start to decline. This is suggestive of resilience of *Methanobrevibacter* spp. to the seaweed-induced changes in the microenvironment compared to the *Methanomassilicoccaceae* and would explain the comparatively modest reductions in methanogen abundance reported previously over shorter experimental periods (Roque et al., 2019). *Methanomassilicoccaceae* spp. (formerly Rumen Cluster C) produce CH_4_ via the reduction of methyl groups (Poulsen et al., 2013; Lang et al., 2015) rather than via the hydrogenotrophic pathway employed by *Methanobrevibacter gottschalkii* and *ruminantium* clades, indicating that this pathway is inhibited to a greater extent by *A. taxiformis* bioactives. The disparity in response to seaweed supplementation among methanogen species is likely multifaceted. Bromoform is the principle anti-methanogenic metabolite found in *Asparagopsis* species (Paul et al., 2006) and blocks the transfer of methyl groups as well as serving as an alternative electron accepter (Patra et al., 2017), which perhaps most readily explains the rapid decline of methylotrophic species observed here. Further, a change in bacterial composition would result in changes to the substrate profile available to the methanogens resulting from the shifts in bacterial composition would indirectly influence archaeal metabolism, as would the changes in the fermenter microenvironment (e.g., H_2_ & CO_2_ levels), and these factors may not affect all methanogenic species in the same manner. Moreover, seaweeds including *A. taxiformis* possess a multitude of other secondary metabolites (e.g. phlorotannins) which are known to impact microbial communities (Ku-Vera et al., 2020) and these may have directly or indirectly contributed to methanogen dynamics in the present study. However given that the other seaweed species tested here didn’t induce major changes in microbial composition, this may not be the case. Individual methanogens could also vary in their resistance to inhibitory compounds, and our data may simply reflect greater resilience of the *Methanobrevibacter* species to the deleterious effects of seaweed bioactives (Ungerfeld et al., 2004). Protozoa-associated methanogens contribute up to 25% of ruminal CH_4_ (Newbold et al., 1995), and numbers of protozoa typically decline over time in the RUSITEC system regardless of treatment (Lengowski et al., 2016; Mateos et al., 2017). While we did not assess the protozoan community in this study, it is likely that a decline in protozoan abundance over time would have at least partially contributed to the reduction in overall methanogen abundance observed here. 16S rRNA data offer limited mechanistic insight, and future studies using shotgun metagenomics or metatranscriptomics may provide clarity as to the mechanisms underpinning these observations.

The fermentation data indicated that the reduction in methanogenesis and inhibition of the methanogenic community was accompanied by a general depression in microbial activity measured by a decline in VFA production and fibre degradation (Terry et al., 2022). This was mirrored in a substantial change in the bacteriome composition, as evidenced by diversity and differential abundance analyses here, one which became more pronounced with advancing time, and was related to important performance metrics like CH_4_ and VFA production. The core microbiome encompasses the most ecologically and functionally important taxa in an environment under given sampling conditions (Neu et al., 2021), and therefore any treatment that disrupts this group may have negative implications for the ecosystem as a whole. In addition to strong relationships between core bacterial genera and fermentation variables for the *A. taxiformis* samples, 19 of the 32 core bacterial genera were DA between *A. taxiformis* and control samples during at least one phase of the study. This suggests that there may be a perturbation of the core microbiome which may have negative effects for essential rumen functions. This was borne out by the reduction in abundances of established bacteria including prominent fibre degraders and VFA producers throughout the experiment suggesting a potential negative impact on animal performance.

A member of the core microbiome, *Prevotella* is a routinely reported as the most abundant rumen microbial genus, prominent in carbohydrate and nitrogen metabolism (Kim et al., 2017). *A. taxiformis* increased the abundance of *Prevotella* during the intermediate phase of the experiment, while differences in the stable phase did not reach significance. Several poorly characterized genera of *Prevotellaceae* were also increased by *A. taxiformis*. It has been speculated that *Prevotella* spp. may redirect excess H_2_ to propionate when CH_4_ is inhibited in the rumen (Aguilar-Marin et al., 2020). However, as propionate levels were reduced by *A. taxiformis* supplementation, increases in *Prevotella* here more likely reflect niche transition of various *Prevotella* species as the abundances of several other established rumen microbes declined.

Acetate, propionate and butyrate are the major VFA associated with ruminal metabolism (Bergman, 1990). In our companion study, *A. taxiformis* reduced total VFA throughout the experiment and the molar proportions of acetate and propionate in the latter phases, although the proportion of butyrate was higher (Terry et al., 2022). While it is not reliable to directly extrapolate from *in vitro* findings, such changes may have negative impacts on animal performance given the importance of propionate and acetate to gluconeogenesis and fatty acid synthesis in the host, respectively (Aschenbach et al., 2010; Bauman et al., 1970). A major pathway for the production of ruminal propionate is via interactions between succinate-producing and -utilizing species (Hobson and Stewart, 1997). *Selenomonas ruminantium* uses succinate in its role as a principal propionate producer in the rumen (Sawanon & Kobayashi, 2006), and A. *taxiformis* reduced the abundance of *Selenomonas*. The abundances of other succinate-utilizers were also reduced, including *Schwartzia* (van Gylswyk et al., 1997) and *Succiniclastium* (van Gylswyk, 1995), indicating that the inhibition of succinate-utilizing bacteria is likely responsible for the reduction in propionate with *A. taxiformis*. However, the abundance of genera containing known succinate producers including *Succinivibrio* and *Ruminobacter* were also increased with *A. taxiformis* samples, suggesting alternative roles for ruminal succinate beyond propionate production during *A. taxiformis* supplementation.

A primary role of the rumen bacteria is the degradation of recalcitrant lignocellulosic biomass by a number of specialized fibre-degrading bacteria, principally *Fibrobacter succinogenes* (Ransom-Jones et al., 2012). Furthermore, fibre degradation is a major source of acetate in the rumen (Hua et al., 2017). Our companion study showed reduced fibre degradation within the *A. taxiformis* samples during the adaption phase though it had recovered by the stable phase, and this was reflected by the significantly lower abundance of *Fibrobacter, Ruminococcus*, and other less common cellulolytic genera like *Ruminclostridium* (Karri et al., 2021; Ren et al., 2019) in *A. taxiformis* samples compared to the controls. The abundance of *Fibrobacter* was also strongly positively correlated with CH_4_ levels associated with *A. taxiformis*, and correspondingly negatively correlated with H_2_ levels. While fibre degradation was only impacted in the adaptation phase (Terry et al., 2022) the abundance of *Fibrobacter* was lower in the controls throughout the experiment, and *Ruminococcus* in the stable phase, suggesting that other microbial groups may have filled the niche for fibre degradation in the latter stages of the experiment, though it should be noted that microbial abundance and activity do not necessarily correlate in complex microbial ecosystem (Hunt et al., 2013). Multiple members of the *Lachnospiraceae* family possess cellulolytic capabilities and are capable of butyrate synthesis (Meehan & Beiko, 2014). *Rosburia* is a prominent butyrate producer (Barcenilla et al., 2000) but was inhibited by *A. taxiformis* here; even though butyrate was the only major VFA to increase in molar proportion during supplementation. Therefore, we speculate that the increased abundance of several poorly described *Lachnospiraceae* genera associated with *A. taxiformis* (e.g. *Lachnospiraceae AC2004, Lachnospiraceae FCS020, Lachnospiraceae NK4A136*) may be responsible for both maintaining fibre degradation in the face of reduced abundance of cellulolytic bacterium and may also have contributed to elevated butyrate levels. This possibility is more likely as no significant effect was observed on the abundance of *Butyrivibrio*, the predominant butyrate producer in the rumen (Moon et al., 2008).

We also noted significantly higher abundances of *Bifidobacterium* in the *A. taxiformis* samples. *Bifidobacterium* is known for its probiotic properties and can adapt to a wide range of substrates (Pokusaeva et al., 2011). Its role in the rumen of adult cattle is less well defined, but elevated abundance is associated with improved feed efficiency (Abe et al., 1995; McLoughlin et al., 2020), and this may be an indicator of rumen microbial adaptation to *A. taxiformis*.

The anti-methanogenic effect of *Asparagopsis* species is attributed to their high bromoform content (Machado et al., 2018). Chemical analysis indicated that only *A. taxiformis* contained measurable amounts of bromoform, as neither *P. mollis n*or *M. japonica* possessed had any detectable amounts, likely explaining their negligible impact on methanogenesis and individual archaeal species. Their lack of impact on the microbiomes generally was surprising, as other seaweeds have been shown to impart significant impact on rumen microbes (Künzel et al., 2022; Zhou et al., 2018) due to the wide range of bioactives present in algae. However, we noted only small impacts on alpha diversity metrics, and shifts in the abundance of minor members of the microbiome. While unlikely to have any potential as an anti-methanogenic feed additive, future studies may examine different doses of *P. mollis* and *M. japonica* to investigate potential prebiotic effects of these algae on the rumen and its microbes.

In summary, this study evaluated the impact of three red seaweeds on the bacterial and archaeal communities in a simulated rumen environment over a 13-day period. The novel red algae *P. mollis* and *M. japonica* had no measureable impact on the microbiome. We found that the inhibition of methanogenesis following *A. taxiformis* supplementation reported in our companion study (Terry et al., 2022) was mirrored by substantial shifts in the microbiome. The rumen methanogens experienced a near-total collapse following *A. taxiformis* supplementation, with few archaeal reads recovered during the stable phase. Similarly, the suppression in VFA synthesis by *A. taxiformis* was underpinned by inhibition of many taxa involved in acetate and propionate synthesis, as well as fibre degradation. *A. taxiformis* is receiving enormous attention for its role as an anti-methanogenic feed supplement in ruminants and has recently been commercialized, and these data provide the first prolonged exploration of rumen microbial dynamics in response to *A. taxiformis* feeding. In-depth studies using shotgun metagenomics or metatranscriptomics may provide a more comprehensive understanding of ruminal microbial dynamics following seaweed feeding, particularly in allowing the simultaneous evaluation of all prokaryotic and eukaryotic communities of the rumen.

## 8. Author Contributions

TAM, DWA, KAB, RJG, and ST designed the experiment. PM, ST and RJG performed the laboratory work. EOH and RJG performed bioinformatics and data analysis. EOH wrote the manuscript with revisions provided by all authors. RJG and KAB supervised the experiment and provided resources.

## 10. Acknowledgments

We thank the platform personnel at the Centre d’expertise et de services Génome Québec for their expertise in performing amplicon sequencing.

## 12. Data Availability Statement

All sequencing data included in this manuscript has been deposited in the Short Read Archive under accession number SUB11925040

